# How Ant Genomes Repeatedly Reinvent Venom

**DOI:** 10.64898/2026.02.12.705515

**Authors:** Frank Andreas Weitz, Francisco Hita Garcia, Bjoern Marcus von Reumont, Burkhard Rost, Ivan Koludarov

## Abstract

**Background:** Venoms from ants (*Formicidae*) are chemically extraordinarily diverse, yet their genomic architecture and evolutionary dynamics remain opaque. The “*aculeatoxin hypothesis*” assumes that most venom peptides in stinging insects (*aculeates*) evolutionarily derive from a single gene family, the aculeatoxins. The recent refutation of this hypothesis for bees raises fundamental questions about venom evolution in ants, which contribute most aculeatoxins.

**Methods:** In order to trace venom peptide evolution, we performed synteny-aware comparative genomics across 25 ant species spanning major subfamilies. We analyzed phylogeny through sequences, structures, and embeddings from Protein Language Models (pLMs).

**Results:** We identified three conserved genomic regions (GR1-GR3) as evolutionary hotspots for ant venom genes, each exhibiting distinct evolutionary dynamics. Most remarkably, we discovered genuine melittin orthologs in ants at the conserved bee syntenic position (GR2), pushing the origin of this scaffold back to early aculeates, with a possible origin deeper in Hymenoptera. Gene copy numbers vary dramatically (0-17 genes per region), with predatory species showing expansions and *formicine* ants (subfamily Formicinae, which rely on formic acid spraying) showing reductions. Twenty-two distinct toxin clades emerge, with region-specific distributions suggesting repeated recruitment to conserved platforms.

**Significance:** Ants evolutionarily succeed by combining single-copy conservation (bee-like at GR2), massive gene duplication (snake-like at GR1), and repeated lineage-specific recruitment to conserved genomic platforms (GR3). This multi-modal evolution on stable genomic scaffolds, evidenced here, reconciles previously conflicting models of venom evolution and reveals how genomic architecture constrains, yet repeatedly enables, molecular innovation. Our findings highlight a general principle of genome evolution: complex adaptive traits can arise not from a single origin or mechanism, but through recurrent reuse of permissive genomic loci shaped by ecology. Such principles are likely relevant not only to other venomous animals, but more broadly to the evolution of complex multi-genic traits.

## Introduction

### Venom evolution at the crossroads

Venom systems represent one of nature’s most successful innovations, evolving independently over 100 times across the animal kingdom ^1^. Two contrasting paradigms have emerged as explanations. The “snake model” explains venom evolution as a **duplication-driven process**, in which repeated gene duplication, positive selection, and birth-and-death dynamics generate expansive toxin gene families ^2^. In contrast, the “wasps and bees model,” emerging from recent genomic studies, reveals venoms largely encoded by **single-copy genes retained at conserved syntenic loci**, where **regulatory co-option rather than gene duplication** underlies venom diversification ^3,4^. Where do ants fit in this dichotomy? With over 13,800 species exhibiting extraordinary venom diversity, from protein-rich cocktails to alkaloid-dominated secretions to formic acid sprays, ants potentially hold the key to understanding how different evolutionary strategies can operate within a single lineage ^5–7^.

### Aculeatoxin controversy and ant centrality

The influential “aculeatoxin hypothesis” proposed that all short aculeate venom peptides derive from a single ancestral superfamily, based primarily on similarities between signal and propeptide sequences ^8^. Since *Formicidae* (ant) sequences constitute about 70% of proposed members of aculeatoxins family, testing this hypothesis requires comprehensive genomic analysis across *Formicidae*.

Our recent work on 32 bee genomes found no support for aculeatoxins: bee toxins occupy lineage-specific loci and cluster by taxonomy rather than function in protein embedding space ^4^. However, that study included limited ant genomes, one showing a melittin-like gene from Asian clonal ant *Vollenhovia emeryi* at the conserved position but with a highly divergent mature sequence, possibly even a pseudogene. The single published *venomics*-oriented ant genome (*Tetramorium bicarinatum*) revealed tandem toxin arrays but lacked comparative context ^9^.

Here, we present the first comprehensive genomic analysis for the evolution of ant toxic peptides, examining 25 species across all major subfamilies. We integrate: (1) microsynteny analysis to establish orthology and trace genomic architecture; (2) phylogenetics to resolve evolutionary relationships; and (3) protein embeddings to detect remote homology and convergence patterns. This multi-faceted approach reveals how ants employ multiple evolutionary strategies simultaneously on conserved genomic platforms.

## Results and Discussion

### Three conserved regions suggest three different evolutionary strategies

Our synteny-aware analysis revealed three main genomic regions (dubbed GR1-GR3, Figures 1, 2 and 3) repeatedly harboring ant venom genes, each exhibiting fundamentally different evolutionary dynamics that illuminate the flexibility of venom evolution within a single lineage.

**Figure 1.**
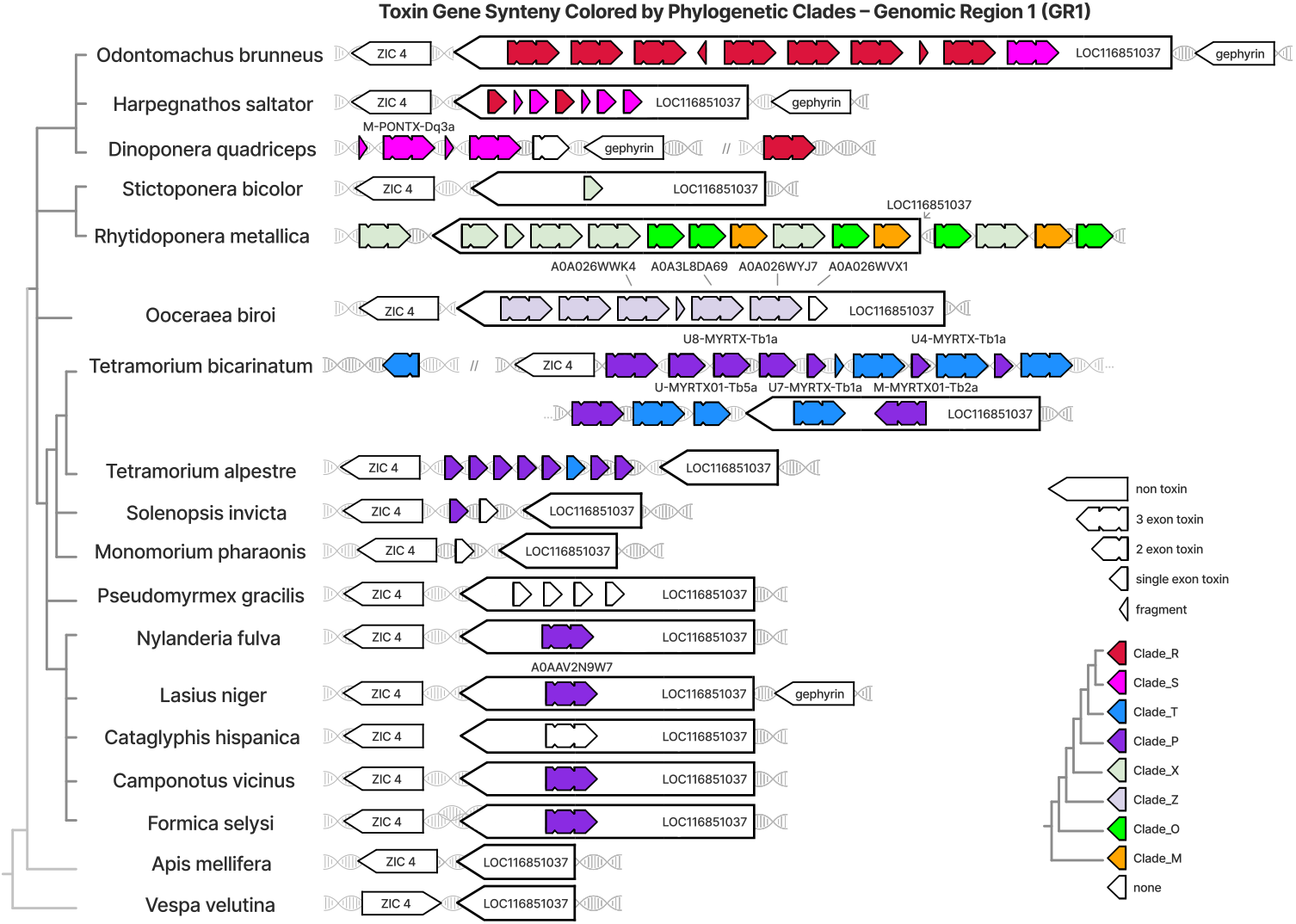
Genomic Region 1 (GR1) shows duplication-driven birth-and-death dynamics. GR1 is a recurrent venom locus displaying extensive copy-number variation across ant genomes (0–17 toxin loci per species). Gene models at this region span intact multi-exon genes, single-exon derivatives, and truncated fragments/pseudogenes, consistent with duplication and loss. Arrowheads show orientation and internal notches mark exon–exon junctions (two notches → three exons; one notch → two exons; no notches → single-exon). The drawn horizontal lengths are purely schematic for clarity (not to scale) and should not be interpreted as quantitative measures of coding-sequence length; the narrow triangular shape denotes a truncated fragment, while the long arrows containing gene names denotes a representative full-length non-toxin construct.

**Figure 2.**
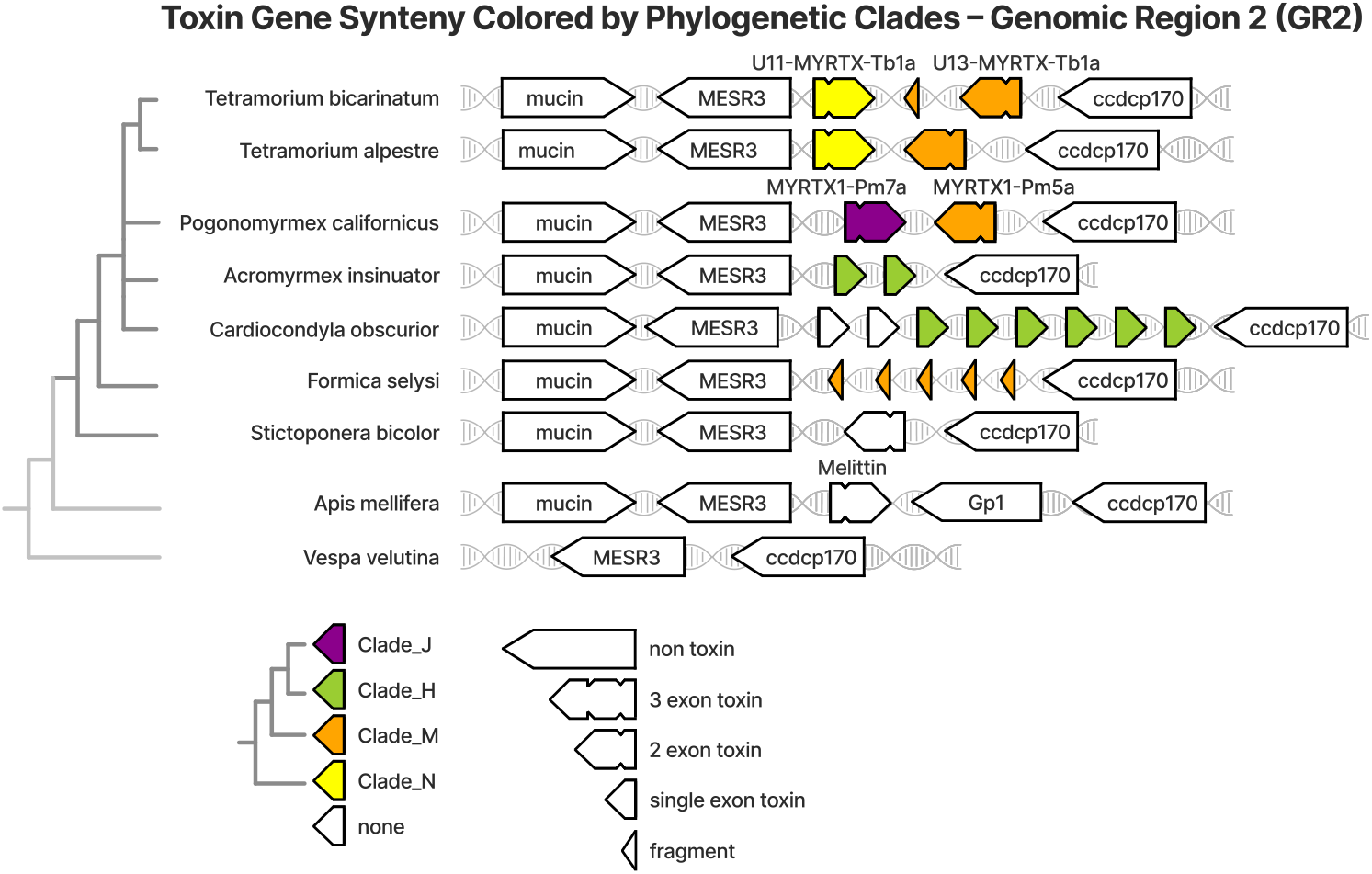
Genomic Region 2. Deeply conserved melittin locus. GR2 occupies the MESR3–CCDC-p170 syntenic interval and in multiple ant lineages carries two-exon venom genes with the canonical organization (exon 1 = signal/propeptide; exon 2 = mature peptide), exemplified by U11-MYRTX-Tb1a and U13-MYRTX-Tb1a in Tetramorium and orthologs in Pogonomyrmex. Positional and structural conservation, together with functional evidence for at least one ant paralog (U11-MYRTX-Tb1a causes paralysis in insects, presumably via K^+^-channel modulation or by permeabilizing membranes ^12,13^), supports orthology with bee melittin; this implies a melittin-like scaffold present in the aculeate ancestor and differentially retained, simplified (single-exon variants) or lost across lineages. Arrowheads show orientation and internal notches mark exon–exon junctions (two notches → three exons; one notch → two exons; no notches → single-exon). The drawn horizontal lengths are purely schematic for clarity (not to scale) and should not be interpreted as quantitative measures of coding-sequence length; the narrow triangular shape denotes a truncated fragment, while the long arrows containing gene names denotes a representative full-length non-toxin construct. GR2 is conserved in these ants, but we could not locate any toxin genes in: Odontomachus brunneus, Trachymyrmex zeteki, Solenopsis invicta, Pseudomyrmex gracilis, Pogonomyrmex barbatus, Ooceraea biroi, Nylanderia fulva, Monomorium pharaonis, Linepithema humile, Harpegnathos saltator, Dinoponera quadriceps, Camponotus vicinus, Paraponera clavata, Leptanilla sp.

**Figure 3.**
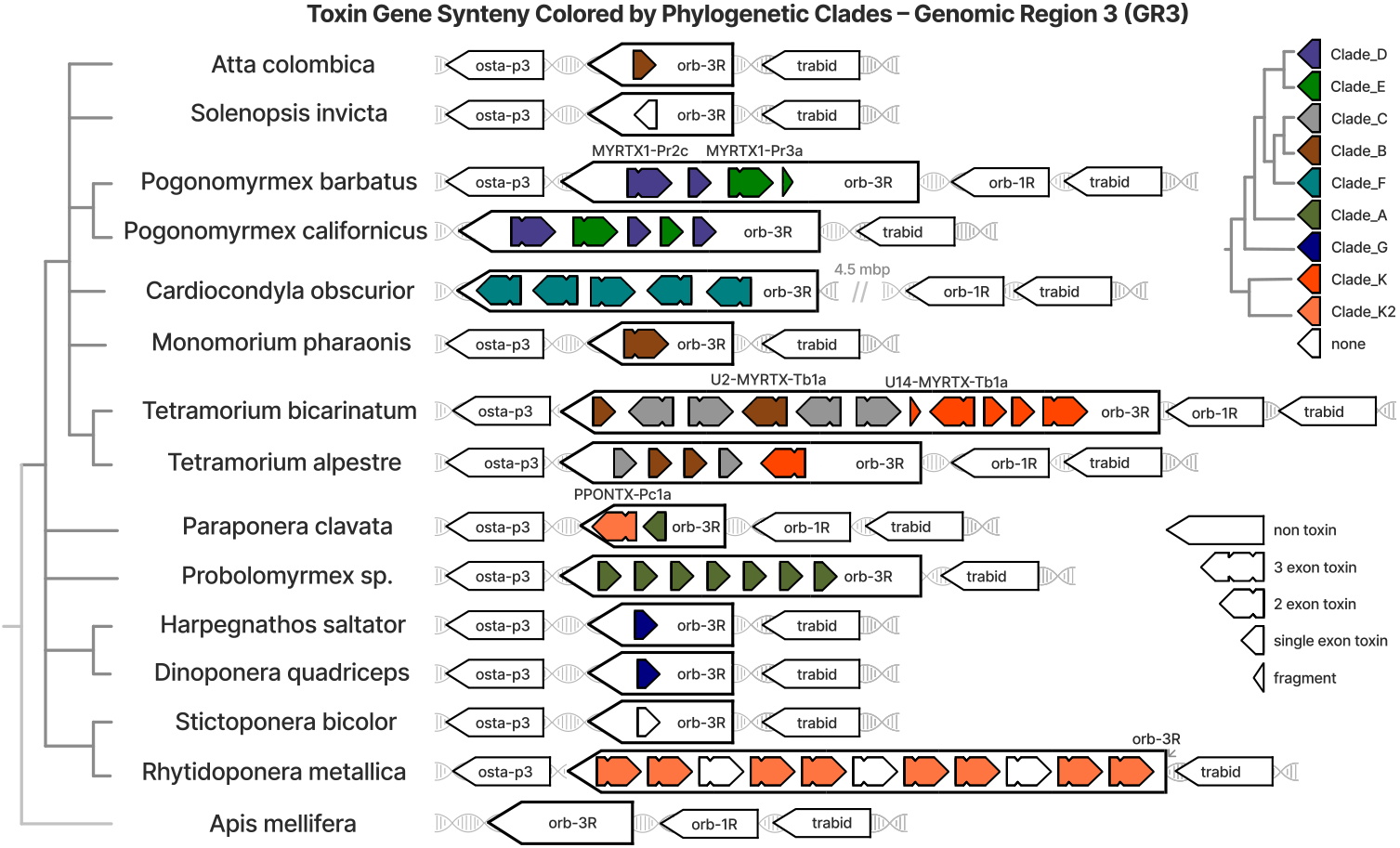
Genomic Region 3. Repeated recruitment loci. GR3 is a stable syntenic scaffold repeatedly colonized by different toxin families in different lineages. Poneroids deploy poneratoxins here (e.g., the NaV-modulating PC-PONTX-Pc1a), myrmicines recruit MYRTX toxins and lineage-specific radiations, and Cardiocondyla contains a unique five-gene tandem array of Clade F toxins. The repeated occupation of the same genomic address by phylogenetically distinct toxin families, together with mixed gene architectures (multi-exon, single-exon, fragments), indicates recurrent recruitment coupled with birth-and-death turnover acting on a taxonomically diverse substrate. Arrowheads show orientation and internal notches mark exon–exon junctions (two notches → three exons; one notch → two exons; no notches → single-exon). The drawn horizontal lengths are purely schematic for clarity (not to scale) and should not be interpreted as quantitative measures of coding-sequence length; the narrow triangular shape denotes a truncated fragment, while the long arrows containing gene names denotes a representative full-length non-toxin construct. GR3 is conserved in these ants, but we could not locate any toxin genes in: Atta colombica, Ooceraea biroi, Pseudomyrmex gracilis, Nylanderia fulva, Lasius niger, Cataglyphis hispanica, Camponotus vicinus, Formica selysi, Mystrium camillae.

### GR1: Snake-like duplication factory

GR1 exemplifies classical *birth-and-death* evolution in which repeated gene duplication is coupled with frequent gene loss and pseudogenization, resulting in extreme copy-number variation (0-17 loci, Fig. 1). The region shows striking ecological correlation: predatory species with functional stings (*Tetramorium bicarinatum*: 17 copies; *Odontomachus, Dinoponera, Harpegnathos*: 4-8 copies) maintain expanded arrays, while formicine ants that spray formic acid retain single or no copies.

Within expanded genomic gene arrays generated by local duplication, we observe the full spectrum of birth-and-death dynamics: intact multi-exon genes preserving signal peptide and propeptide architectures, partially degraded single-exon variants, and fragmentary pseudogenes. The retention of introns in duplicated copies argues against retroposition (RNA-mediated duplication) and instead supports local DNA-level gene duplication as the primary mechanism of array expansion.

Phylogenetically, GR1 toxins segregate by ant lineage: *poneroid* ants share clades S and R (including the antimicrobial M-PONTX-Dq3a), whereas *myrmicine* ants (subfamily *Myrmicinae*) possess distinct clades T, U, and W. Within the genus *Tetramorium*, both ancient clades shared across the genus, including those retained in *Tetramorium alpestre*, and recent lineage-specific expansions (Clade U) are observed, illustrating ongoing diversification. This pattern identifies *Tetramorium* as a hotspot of recurrent local gene duplication, exhibiting unusually frequent GR1 copy turnover relative to other ant lineages. Such a high copy-turnover might be related to extreme survival pressure ^10^.

### GR2: Melittin locus deeply conserved

The most striking discovery is the presence of two-exon venom genes at the exact syntenic position (MESR3–CCDC-p170 interval) at which bees encode melittin (Fig. 2). These ant sequences, U11-MYRTX-Tb1a and U13-MYRTX-Tb1a in *Tetramorium*, plus orthologs in *Pogonomyrmex*, preserve the standard architecture of aculeate venom peptides: the signal- and pro-peptide in exon 1, and the mature peptide in exon 2.

This conservation of position and gene model, combined with functional evidence (U11-MYRTX-Tb1a causes paralysis in insects, presumably via K^+^ channel modulation or by permeabilizing membranes in a pore-forming manner ^11–13^), strongly suggests orthology with bee melittin. The implications are profound: the melittin scaffold likely existed in the *aculeate* ancestor over 200 million years ago, with lineage-specific sequence divergence obscuring the relation at the sequence level.

Some ants retain two-exon genes, others have single-exon variants or completely lost the gene. This patchy diversity across ants somewhat mirrors the bee pattern where melittin is preserved in most lineages but lost in a few others. This “use it or lose it” pattern at a conserved locus contrasts sharply with the dynamic expansions at GR1.

### GR3: Recruitment platform

GR3 represents a novel evolutionary mode in which a stable syntenic genomic scaffold is repeatedly colonized by distinct toxin families in different ant lineages (Fig. 3). *Poneroid* ants recruit poneratoxins at this locus, including the NaV-modulating PC-PONTX-Pc1a responsible for extremely painful stings. In contrast, *myrmicine* ants (subfamily *Myrmicinae*) independently deploy MYRTX peptides of unknown function (U*-MYRTX) alongside lineage-specific radiations. Notably, the myrmicine species *Cardiocondyla obscurior* exhibits a unique five-gene tandem array of Clade F toxins which, although absent from other taxa, forms a basal clade relative to all other GR3-associated sequences.

This pattern, “same genomic address with different molecular tenants”, suggests that certain chromosomal contexts are inherently permissive for venom gene expression. This might be due to regulatory landscapes or to the chromatin architecture. The mosaic of multi-exon genes, single-exon variants, and fragments within GR3 again supports *birth-and-death dynamics* which, in contrast to the situation for GR1, operates on taxonomically diverse substrates.

### Aculeatoxins: 22 clades tell 22 stories

Our phylogenetic analysis recovered 22 distinct toxin clades with region-biased distributions. Crucially, these clades segregate by ant taxonomy (subfamily or tribe) rather than forming a single pan-aculeate superfamily. Together with syntenic evidence showing that different toxin clades occupy distinct genomic loci, this pattern is inconsistent with the aculeatoxin hypothesis, which posits a monophyletic origin of all aculeate toxins from a single ancestral gene at the stem *Aculeata*. Instead, our results support a polyphyletic origin of aculeate venom toxins, involving independent recruitment or diversification events within different aculeate lineages. This does not preclude deeper homology among venom genes at the level of stem *Hymenoptera*, but it rejects a single shared aculeate-specific toxin ancestor.

Embeddings from protein Language Models (pLMs) provide orthogonal support: when visualized in UMAP space, ant toxins cluster by genomic region and taxonomic origin, not by functional similarity (Fig. 4). Even when including signal peptides (the primary evidence for classification as *aculeatoxins*), sequences separate by lineage. This pattern, also seen in our bee analysis ^4^, indicates convergent evolution of similar functions rather than divergence from a common ancestral toxin.

**Figure 4.**
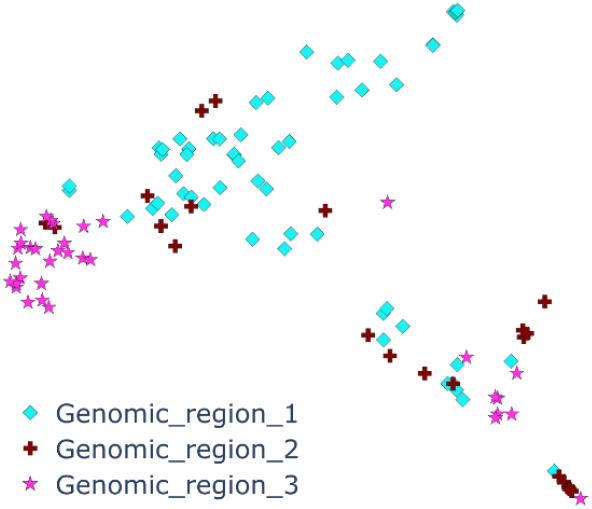
ProtT5 Embeddings show disperse clustering, with sequences from the same genomic region favoring proximity. Protein-sequence embeddings (including signal peptides; see Methods) projected into two-dimensional UMAP space. Points are colored by Genomic Region. Embeddings group by lineage and genomic region rather than by inferred function, supporting convergent recruitment of toxin-like sequences rather than a single ancestral “aculeatoxin” family.

### Ecological drivers and functional correlations

The correlation between venom complexity and ecology is striking. Predatory ants with functional stings consistently show expanded repertoires at GR1 and GR3. *Formicine* ants, which secondarily lost the ability to sting and evolved formic acid spraying, show minimal peptide toxins, often just single genes or complete absence at these loci.

This pattern extends to specific clades: poneratoxin-related sequences (targeting vertebrate pain pathways) occur in large predatory species that use stings defensively against vertebrates. *Myrmicine*-specific expansions may target insect prey or microbes. The *Cardiocondyla*-specific radiation at GR3 could reflect unique social or ecological pressures in this genus.

### Synthesis: ants reveal the full complexity of venom evolution

Our findings position ants as uniquely revealing the full complexity of venom evolution across *Hymenoptera*. Five themes emerge.

1. **Multiple evolutionary mechanisms coexist**. Unlike bees, which primarily rely on regulatory co-option of single-copy genes, or snakes, which largely diversify venoms through extensive gene duplication, ants employ both duplication-driven expansion and regulatory recruitment, alongside a lineage-specific locus reuse mechanism in which different toxin families independently colonize conserved syntenic regions. Possibly the number of different evolutionary mechanisms provides a better proxy to the complexity of ants than the mere number of lineages or species (roughly similar between ants and bees).
2. **Mosaic patterns reveal deep but uneven homology**. The discovery of the melittin locus in ants demonstrates that some venom positions are genuinely orthologous across *aculeates*, preserved for over 200 million years despite divergence beyond recognition by sequence similarity alone. In contrast, other venom loci show clear lineage-specific origins.
3. **Genomic architecture channels innovation**. The recurrent involvement of GR1–GR3 across ant evolution suggests that these conserved syntenic loci possess intrinsic properties that facilitate venom gene expression and turnover. These could include regulatory context, chromatin accessibility, or nuclear positioning.
4. **Ancient loci are differentially retained**. Functional toxin genes in one lineage correspond to pseudogenes in others (e.g. melittin, mastoparan, and poneratoxin loci ^4^). This observation indicates ancient genomic presence followed by lineage-specific retention or loss.
5. **Ecological specialization drives genomic complexity**. The strong interplay between predatory venom use, such as prey capture, immobilization, and pain induction, and toxin gene expansion suggests that ecological reliance on venom actively promotes the retention, diversification, and amplification of duplicated genes, rather than duplication being selectively neutral.

Overall, our results are incompatible with both models of entirely independent *de novo* origins of venom systems and with a single monophyletic *aculeate* toxin ancestry. Instead, advanced sequence analysis enabled by protein language models and comparative genomics reveals a dynamic evolutionary landscape characterized by deep homology, repeated recruitment, and ongoing innovation in aculeate venom systems.

## Conclusion

Ant venom evolution operates through multiple mechanisms on three conserved genomic platforms. Our discovery of melittin orthologs at the bee syntenic position established a deep conservation of genomic architecture despite extreme sequence divergence. The three regions show three modes: massive duplication (GR1), single-copy conservation (GR2), and platform recruitment (GR3). These findings reconcile conflicting models of venom evolution, showing how different strategies can operate simultaneously within a single lineage depending on ecological pressures and genomic context. Combined with evidence for convergent evolution on conserved scaffolds this establishes a new paradigm: venom evolution is channeled by genomic architecture but diversified by ecological selection. Such a multi-modal evolutionary framework likely extends beyond ants. The three principles revealed here - conserved platforms, lineage-specific recruitment, ecology-driven expansion - may explain venom diversity across the tree of life.

## Methods

### Data acquisition

#### Genome assemblies

We retrieved publicly available genome assemblies for the species included in this study from NCBI (accession numbers and assembly versions are given in Supplementary Table 1). Assemblies were used as provided (no additional scaffolding or reassembly) and were chosen to prioritize high contiguity and available annotation where possible. ^14–28^

#### Toxin sequences

We assembled a reference set of known peptide toxin sequences by downloading reviewed and non-redundant (less than 100% sequence identity) records from UniProt ^29^ and by manually extracting additional sequences reported in the literature (mass-spectrometry, transcriptomics, and genomic toxin descriptions). The combined data set (UniProt + literature) served as the query database for all subsequent genome searches.

### Genome searches with MMseqs2

We searched each genome for toxin query and homologs of the toxin query set using MMseqs2 ^30^. Protein searches were performed using translated search modes when necessary to accommodate exon/intron structure in draft assemblies. MMseqs2 was run with high sensitivity to maximize recovery of divergent toxin homologs (sensitivity = 9.5) and an e-value cutoff of 1e-3 for initial hit detection.

### Manual curation and exon boundaries (UGENE + BDGP splice site prediction)

MMseqs2 hits were imported into UGENE ^31^ for manual inspection. For each candidate locus we applied UGENE’s built-in Smith–Waterman algorithm to confirm alignment quality and to detect whether multiple adjacent peptide matches corresponded to separate exons. Short, isolated matches (very small peptide fragments recurring at multiple genomic locations) were considered likely repeats and discarded. Where adjacent hits from the same query occurred on the same scaffold, these were treated as putative exon pairs and inspected for canonical splice signals.

To define exon boundaries, we combined automated splice site prediction available at fruitfly.org ^32^ with manual sequence matching. Predicted donor/acceptor sites were used as a guide and were accepted if (1) they mostly aligned with UGENE-identified exon boundaries via Smith–Waterman alignment, and (2) the resulting exon concatenation produced a coherent open reading frame consistent with the query toxin. In ambiguous cases we manually adjusted exon start/stop positions to preserve reading frame and expected signal/propeptide features. Percent-identity thresholds used during manual verification were relaxed relative to standard protein searches (typically 25–35% depending on query length) to accommodate rapid toxin evolution while limiting spurious matches.

### Extraction of flanking genes and orthology searches

For each confirmed toxin locus we recorded the immediate upstream and downstream protein-coding genes (flanking genes) based on the genome annotation. Coding sequences of these flanking genes were extracted separately and used as queries in a second round of MMseqs2 searches against the full set of genomes. The aim was to (1) locate homologous genomic neighborhoods in other assemblies and (2) distinguish orthologous toxin insertions from lineage-specific insertions or assembly artifacts.

Each flanking CDS was searched independently using MMseqs2 with parameters matching the toxin searches (sensitivity = 9.5; e-value ≤ 1e-3). Hits were compiled and inspected visually to determine which scaffolds carried the conserved gene order expected for orthology. In regions where both flanking genes were recovered but no annotated toxin was present, we re-examined the scaffold with exon-level Smith–Waterman searches (UGENE) to discover highly divergent toxin orthologs.

### Synteny analysis and plotting

From the manual curation of flanking gene hits we inferred conserved genomic neighborhoods. We recorded the presence/absence and relative order of flanking genes and noted whether an identifiable toxin CDS co-occurred at the locus. Using these data we produced synteny figures representing the four principal toxin loci studied.

### Multiple sequence alignment and outlier filtering

Toxin amino-acid sequences were aligned with MAFFT ^33^ using the L-INS-i style parameters appropriate for short, divergent peptides: --localpair --maxiterate 1000. Alignments were inspected visually and a small number of clear outliers (typically ∼2–3 sequences per alignment) that degraded alignment quality were removed. The alignment was then recomputed with the same MAFFT parameters to obtain the final alignment used for phylogenetic inference.

### Phylogenetic inference

We inferred maximum-likelihood trees from the MAFFT alignment using IQ-TREE 2 ^34^. The analysis was run with model selection (MFP) augmented by a fixed set of mixture matrices and branch-rate options, and with intensive support estimation as follows:

iqtree2 -s toxins.fasta -st AA -m MFP -madd LG+C60+F+G -mrate R -bb 10000 -alrt 10000 -bnni -allnni -pers 0.2

Resulting trees were filtered to collapse any node with bootstrap support below 50% to avoid over-interpretation of weakly supported relationships. We then manually inspected the collapsed topology, defined robust ant-specific clades, and annotated these clades for downstream visualization.

### Integration of phylogeny and synteny

Clade assignments from the IQ-TREE analysis were mapped back onto the synteny plots. Each tree-defined ant toxin clade was given a distinct color and these colors were used to mark toxin occurrences in the synteny diagrams to facilitate interpretation of the relationship between sequence evolution and genomic context.

### Sequence embedding and low-dimensional visualization

To provide an orthogonal, alignment-free view of sequence relationships, we generated per-protein embeddings with the ProtT5 ^35^ protein language model. ProtT5 embeddings were computed for all sequences, averaged across residues to produce one fixed-length vector per protein, and subsequently projected to two dimensions using UMAP via the ProtSpace ^36^ workflow. Embedding plots were colored by (1) genomic locus, (2) scaffold/species origin, and (3) phylogenetic clade to aid visual interpretation.

## Abbreviations

CDS: (coding DNA sequence)
GR: (genomic region)
K^+^: (potassium ion)
MYRTX: (Myrmecitoxin, proposed ant venom toxin family)
NaV: (voltage-gated sodium channel)
NCBI: (National Center for Biotechnology Information)
ORF: (open reading frame)
PONTX: (Poneratoxin, proposed ant venom toxin family)
ProtT5: (Protein Transformer T5, protein language model)
UMAP: (Uniform Manifold Approximation and Projection for dimensionality reduction)
UniProt: (Universal Protein Resource)
ccdc-p170: (Coiled-coil domain–containing protein 170).

## Supplementary Material

**Supplementary Figure 1:**
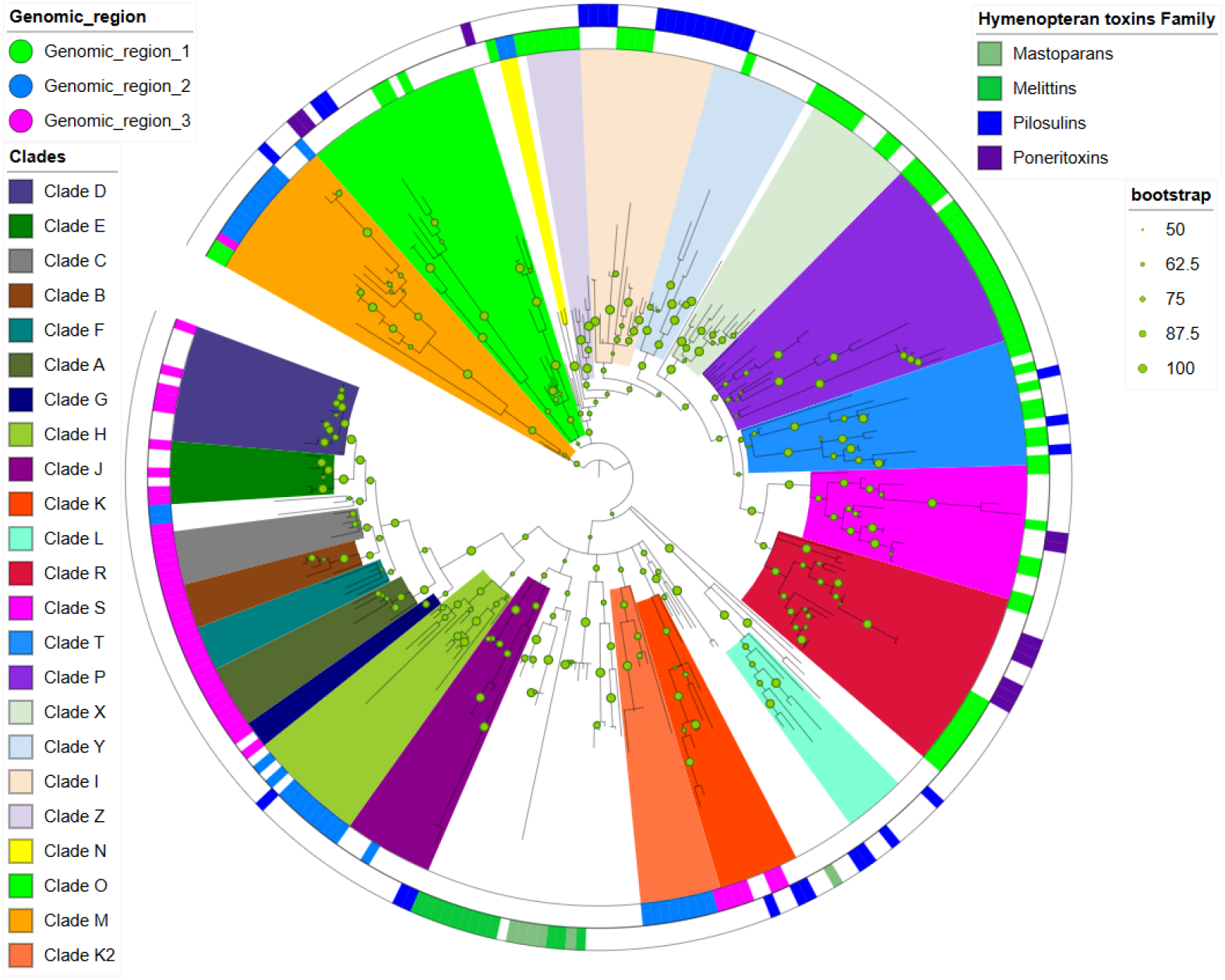
Maximum-likelihood phylogenetic tree of ant toxins together with Hymenopteran toxins such as melittins. Dot sizes next to nodes in the tree indicate bootstrap values as indicated in the legend. The inner annotation ring displays the phylogenetic clades of ant toxins. The middle ring displays the genomic region a toxin is located in. The outer ring contains only the toxins that did not come from the ant toxin analysis directly, but from an unpublished manuscript about Hymenopteran venoms.

**Supplementary Figure 2:**
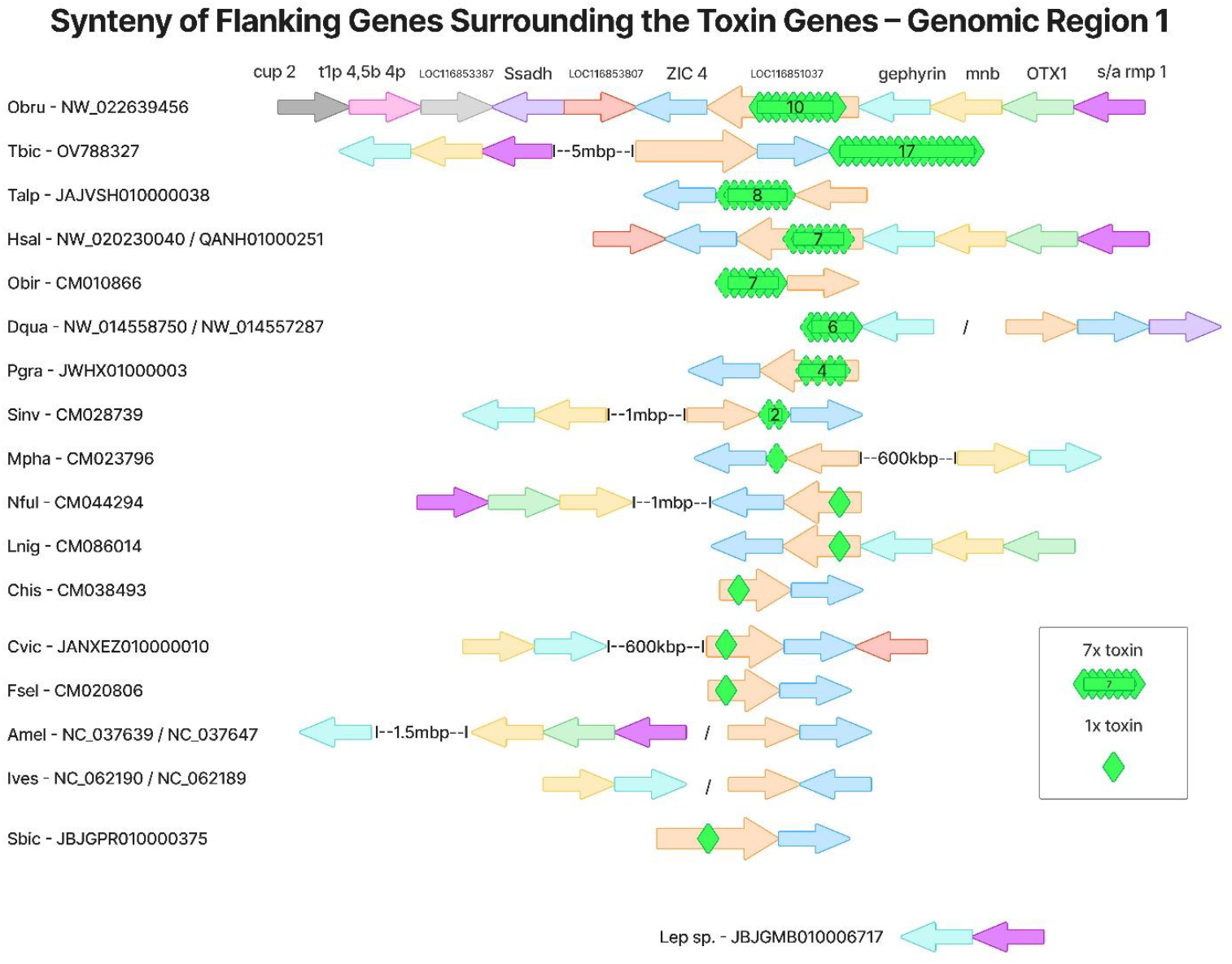
Synteny Figure of GR1 like Figure 1, but with more flanking genes and less detail on toxins. Information on Scaffold / Chromosome are given.

**Supplementary Figure 3:**
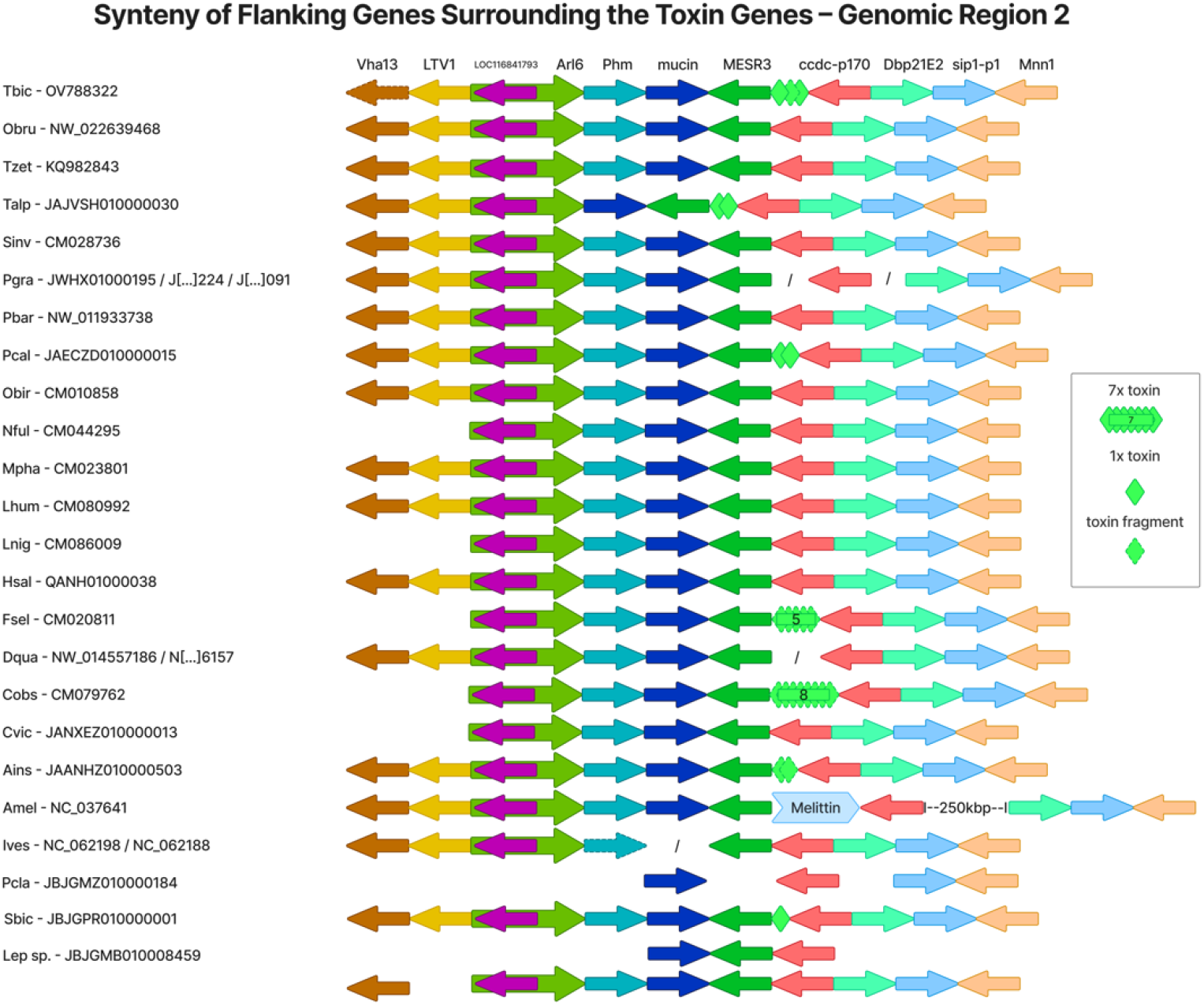
Synteny Figure of GR2 like Figure 2, but with more flanking genes and less detail on toxins. Information on Scaffold / Chromosome are given.

**Supplementary Figure 4:**
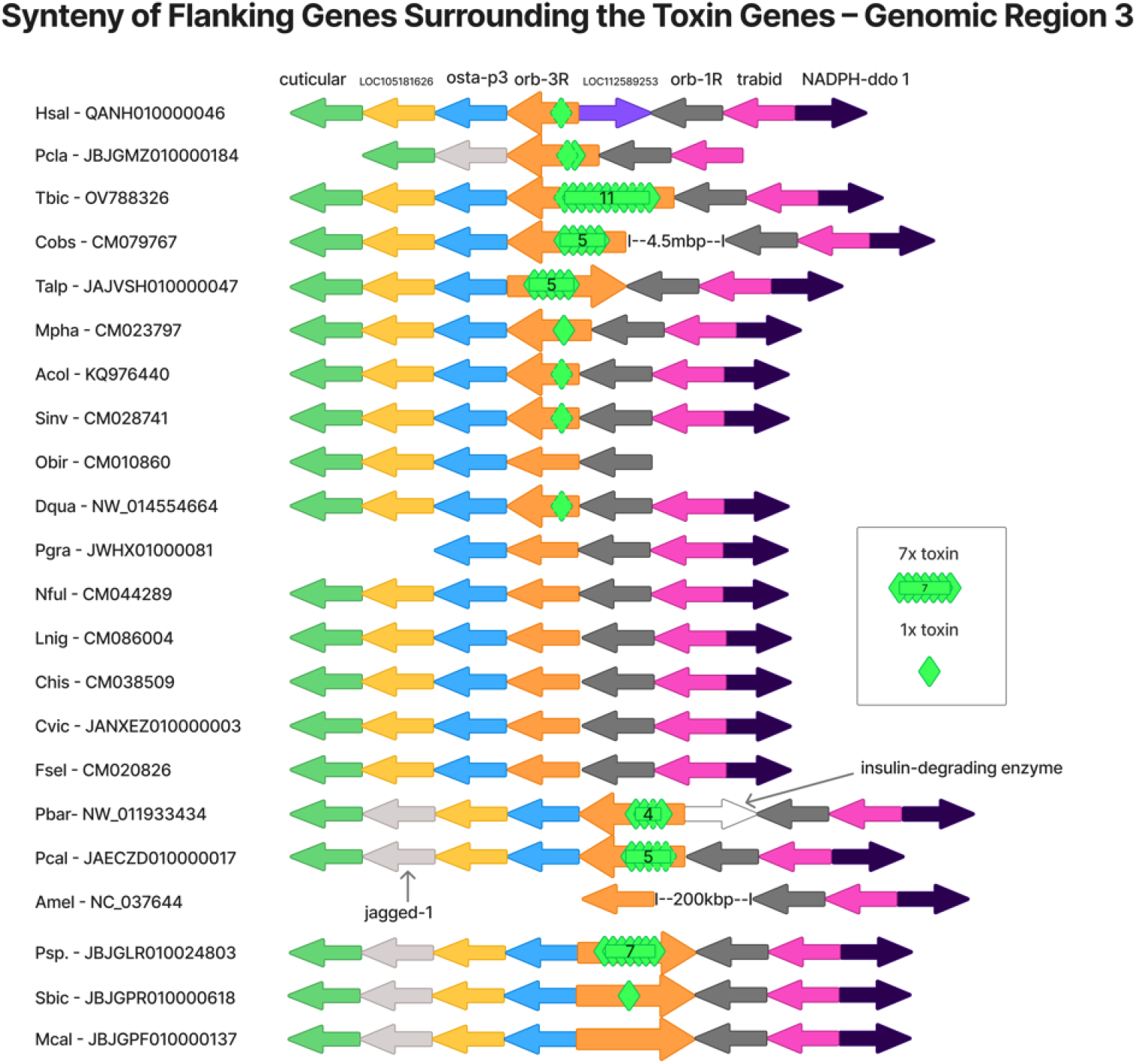
Synteny Figure of GR3 like Figure 3, but with more flanking genes and less detail on toxins. Information on Scaffold / Chromosome are given.

**Supplementary Table 1.**
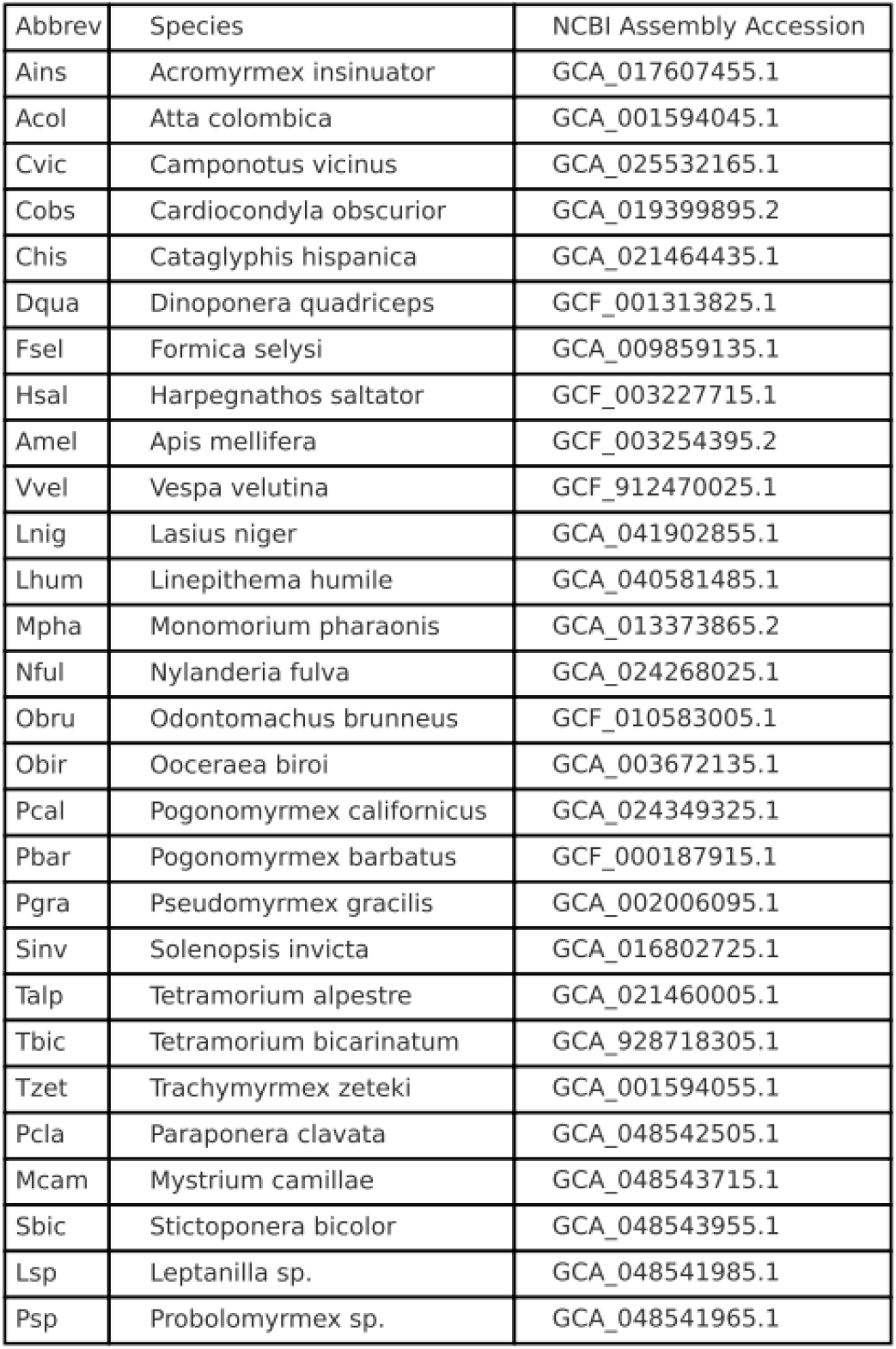
Species codes, full species names, and NCBI assembly accession numbers (including version) for the genomes used in this study.

